# AKIP1 is an inner scaffold component required for centriole integrity

**DOI:** 10.64898/2026.06.03.729831

**Authors:** Akira Sanada, Kei K Ito, Takumi Chinen, Shohei Yamamoto, Masamitsu Fukuyama, Shoji Hata, Daiju Kitagawa

## Abstract

Centrioles are microtubule-based organelles that play crucial roles in various biological processes, including cell division, ciliogenesis and flagellar assembly. These functions require the preservation of centriole structural integrity, yet the molecular mechanisms that maintain this integrity remain incompletely understood. In this study, through large-scale gene co-dependency analysis and ultrastructure expansion microscopy (U-ExM), we identified A-kinase interacting protein 1 (AKIP1) as a novel component of the centriole inner scaffold, a luminal structural framework that supports centriole integrity. Analysis of large-scale functional genomic data from DepMap placed AKIP1 in a functional cluster with the inner scaffold components POC1A, POC1B, and POC5. Consistently, AKIP1 colocalizes with these clustered inner scaffold components in the central core region of centrioles, forming a multicomponent interaction network. We further show that AKIP1 is recruited to centrioles during G2 phase, subsequent to the recruitment of other inner scaffold components, and that its centriolar localization depends on these components. AKIP1 depletion leads to defects in centriolar microtubules and abnormal centriole morphology. Together, our findings establish AKIP1 as a downstream component of the inner scaffold that contributes to the maintenance of centriole integrity.

## Introduction

Centrioles are microtubule-based intracellular structures that are conserved in many eukaryotes^1,2^. In proliferating cells, centrioles recruit pericentriolar material (PCM) to form centrosomes, the major microtubule-organizing centers (MTOCs) that organize intracellular microtubule networks and promote mitotic spindle assembly^3–5^. In quiescent G0 phase cells, centrioles function as basal bodies that template the formation of cilia and flagella^5,6^. These functions require the maintenance of centriole structural integrity, as defects in centriole structure can impair mitotic spindle assembly, ciliogenesis, and flagellar assembly and are associated with ciliopathies and other developmental disorders^7–9^.

In human cells, centrioles are cylindrical structures approximately 250 nm in diameter and 450 nm in length, composed of nine radially arranged microtubule triplets^2,10^. Along their proximal-distal axis, centrioles are divided into three regions: proximal, central, and distal, each of which contains characteristic structures associated with the centriolar microtubule wall^10,11^. In duplicating centrioles, the proximal region contains the cartwheel, which provides the structural foundation for centriole duplication and establishes the ninefold symmetry of the centriole^12,13^. The cartwheel is connected to the microtubule triplets through pinhead structures, while A-C linkers connect adjacent microtubule triplets in the proximal region^11,14^. At the distal end, distal and subdistal appendages are present on the outer surface of the microtubule wall. These appendages mediate centriole docking to the plasma membrane during ciliogenesis and microtubule anchoring during interphase, respectively^10,15^. In the central core region, the inner scaffold supports the cylindrical arrangement of the microtubule wall from the luminal side and is thought to maintain the mechanical strength and long-term stability of centrioles^16–18^.

The molecular architecture of the inner scaffold has been progressively elucidated through cryo-electron tomography and ultrastructure expansion microscopy (U-ExM)^11,17–21^. Cryo-electron tomography analyses of centrioles in *P. tetraurelia* and *C. reinhardtii* have shown that the inner scaffold forms a helical lattice-like structure composed of repeating structural units along the inner surface of the centriolar microtubule wall^18^. These studies further demonstrated that this architecture is evolutionarily conserved^18^. The development of U-ExM, which enables visualization of centriole protein localization at nanoscale resolution, has facilitated the identification of proteins associated with inner scaffold organization, including POC5, POC1A, POC1B, FAM161A, Centrin, WDR90, and CEP350^15,17–20,22^. POC1A forms heterodimers with POC1B and contributes to the organization of a luminal interaction network by recruiting additional inner scaffold components, including POC5^17^. POC5 forms tetramers and associates with Centrin, whereas FAM161A, WDR90, and CEP350 have been implicated in linking the inner scaffold or associated centriolar structures to the microtubule wall^17,18,20,22–26^. In particular, CEP350 promotes the centriolar localization of WDR90, while WDR90 contributes to the connection between the inner scaffold and the centriolar microtubule wall^20,22^. Thus, the inner scaffold is organized by a complex network of interacting proteins, although its full molecular organization remains incompletely defined.

To date, inner scaffold components have been identified largely through candidate-based localization analyses of previously characterized centriolar and ciliary proteins using super-resolution imaging approaches such as U-ExM^11,17,18,20^. However, such candidate-based approaches are unlikely to capture the full repertoire of proteins that build and organize this complex luminal network. In this context, recent analysis of the inner scaffold-associated proteome using proximity-dependent labeling, an unbiased alternative to candidate-based approaches, identified CCDC15 as a novel inner scaffold protein^21^. Unbiased discovery approaches therefore offer complementary routes to identify previously unrecognized inner scaffold components.

As another unbiased, function-oriented approach, large-scale functional genomic datasets from DepMap can be used to identify genes with shared biological roles^27–29^, as already demonstrated in centriole biology by the identification of A-C linker components^14^. Cellular dependence on specific intracellular structures or molecular pathways often varies among cell types, reflecting differences in tissue-specific properties and gene expression programs^27,29^. These variation patterns can be exploited to uncover genes that function in common biological processes^29,30^.

In this study, we performed gene co-dependency analysis using DepMap datasets and identified a functional cluster comprising POC1A, POC1B, and POC5, known components of the centriole inner scaffold, together with AKIP1, which had not previously been characterized as a centriolar protein. AKIP1 localizes to the centriole inner scaffold and appears at centrioles during G2 phase. It associates with the clustered inner scaffold components, and its centriolar localization depends on POC1A and POC5. Loss of AKIP1 causes centriole structural defects, indicating that it contributes to the maintenance of centriole architecture.

## Results

### Identification of AKIP1 as a putative inner scaffold protein through gene co-dependency analysis

Large-scale CRISPR-Cas9 loss-of-function screening datasets from the Cancer Dependency Map (DepMap version:25Q3) project are now publicly available for more than 1,000 cancer cell lines^27,28^. These datasets provide gene dependency profiles that reflect the extent to which each cell line depends on individual genes for proliferation or survival. Because genes involved in the same intracellular structure or molecular pathway often show similar dependency profiles, co-dependency analysis can be used to infer functional relationships among genes^27,29^. To identify novel inner scaffold-associated genes, we applied co-dependency analysis by calculating pairwise Pearson correlations between gene dependency profiles across cell lines (Fig. 1A). Among known inner scaffold components, POC1A, POC1B, and POC5 showed particularly strong co-dependency with one another in this dataset (Table S1A-C). We therefore used these three factors as query genes to search for previously uncharacterized inner scaffold-associated components (Fig. 1B, Table S1D). This analysis identified many centriolar proteins among the top genes co-dependent with POC1A, POC1B, and POC5. These included known inner scaffold components such as CETN2/Centrin-2^18^, as well as factors implicated in the maintenance of centriole structure, including CEP120^31^, CEP44^32^, C2CD3^33^, and HYLS1^34^ (Fig. 1B, C, Table S1D). In addition to these known centriolar genes, A-kinase interacting protein 1 (AKIP1), which had not previously been linked to centriole biology^35^, emerged as the strongest hit (Fig. 1B, Table S1D). We next performed reciprocal gene co-dependency analysis using AKIP1 as the query gene. POC1A, POC1B, and POC5 were among the genes most strongly co-dependent with AKIP1 (Fig. 1D, Table S1E). Furthermore, hierarchical clustering based on gene dependency profiles of AKIP1 and 252 genes annotated as “centriole” or “centrosome” in Gene Ontology placed AKIP1 in the same cluster as POC1A, POC1B, and POC5 (Fig. 1E). Similarly, clustering analysis of known inner scaffold genes showed that POC1A, POC1B, POC5, and AKIP1 formed a distinct cluster, whereas other known inner scaffold components showed weaker correlations with this group (Fig. 1F). Together, these results suggest that AKIP1 is functionally linked to the centriole inner scaffold and is particularly associated with the POC1A-POC1B-POC5 cluster.

**Figure 1.**
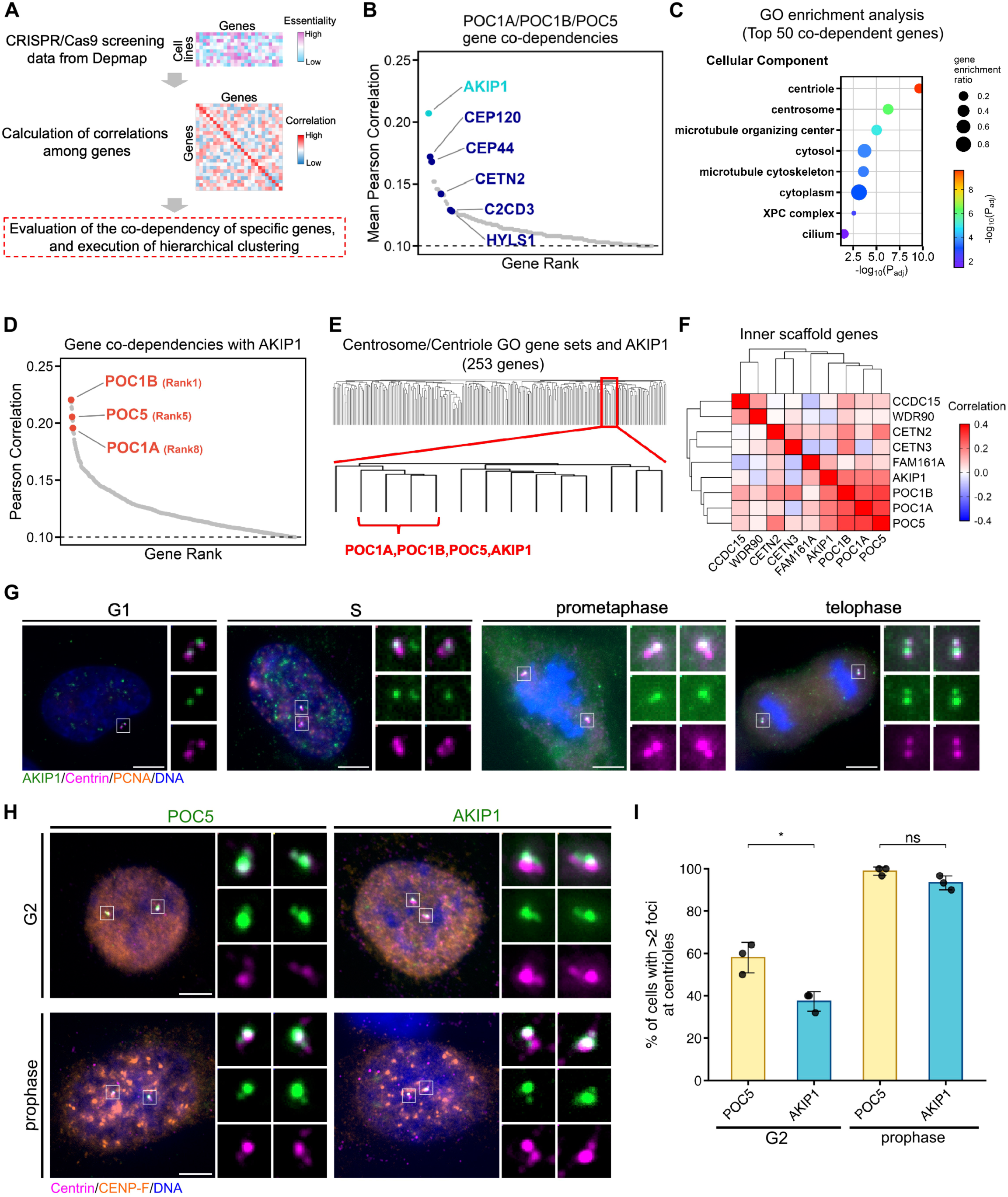
Identification of AKIP1 as a putative inner scaffold protein through gene co-dependency analysis. **(A)** Scheme of gene co-dependency analysis for the exploration of unknown centriole genes. **(B)** Gene co-dependency predictions for POC1A, POC1B, and POC5. Scores represent the average Pearson correlation coefficients of CRISPR gene effect scores between each gene pair and POC1A, POC1B, and POC5. Previously known centriolar structural factors are shown in blue. **(C)** Gene Ontology (GO) enrichment analysis using g:Profiler was conducted for the top 50 genes identified in (B). Enriched GO terms in the Cellular Component category are shown. **(D)** Gene co-dependency predictions for AKIP1. Scores represent the Pearson correlation coefficients of CRISPR gene effect scores between each gene pair. **(E)** Hierarchical clustering of AKIP1 and genes annotated to "centriole" or "centrosome" in the Cellular Component GO terms, using gene correlations. **(F)** Hierarchical clustering of known inner scaffold genes and AKIP1 using gene correlations. **(G)** Localization of endogenous AKIP1 to the centrioles in different cell cycle stages. RPE-1 cells were immunostained with antibodies against AKIP1 (green), Centrin (magenta), and PCNA (orange). DNA was visualized with DAPI. Scale bar, 5 μm. **(H)** Representative immunofluorescence images of RPE-1 cells in G2 phase and prophase. RPE-1 cells were immunostained with antibodies against AKIP1 or POC5 (green), Centrin (magenta), and CENP-F (orange). DNA was visualized with DAPI. Scale bar, 5 μm. **(I)** Bar graphs represent the percentage of cells with more than two AKIP1 or POC5 foci on the centrioles in G2 phase and prophase. Values are the mean percentages ± SD from three independent experiments (n = 50 for G2 phase, n = 30 for prophase from 3 independent experiments). A two-tailed, unpaired Student’s t test was used to obtain the P values. *, P < 0.05; NS, P > 0.05.

### AKIP1 is a component of the centriole inner scaffold

To investigate the relationship between AKIP1 and the inner scaffold, we first examined the localization of AKIP1 in RPE-1 cells by immunofluorescence microscopy. Co-immunostaining for AKIP1 and the centriole marker Centrin revealed that AKIP1 localized to centrioles throughout the cell cycle (Fig. 1G). The number of AKIP1 foci was two until S phase and predominantly four during mitosis (Fig. 1G), suggesting that AKIP1 is recruited to daughter centrioles between S phase and mitosis. Known inner scaffold proteins are incorporated into centrioles at distinct cell-cycle stages: POC1A and POC1B are recruited during S phase^17^, whereas POC5, WDR90, and CCDC15 are incorporated during G2 phase^20,21,23^. To determine when AKIP1 is recruited to centrioles, we identified cells in G2 phase and prophase based on CENP-F localization and nuclear morphology, and then examined AKIP1 localization at their centrioles. AKIP1 was detected at more than two centrioles in approximately 40% of G2-phase cells and in most prophase cells (Fig. 1H, I). Comparison with POC5, which localizes to daughter centrioles from G2 phase, showed that AKIP1 recruitment was less frequent than POC5 recruitment in G2-phase cells (Fig. 1I). These results suggest that AKIP1 is recruited to centrioles after POC5. Together, these observations indicate that AKIP1 is a centriole-localized protein whose recruitment to daughter centrioles begins during G2 phase and is largely established by prophase.

We next examined the subcentriolar localization of AKIP1 using super-resolution microscopy. Ultrastructure expansion microscopy (U-ExM) enables nanoscale mapping of protein localization within centrioles^19^. In both side and top views of RPE-1 centrioles, AKIP1 localized to the central core region and was positioned on the luminal side of centriolar microtubules, similar to POC1A, POC1B, and POC5, which formed a functional cluster with AKIP1, as well as Centrin-2/3, known inner scaffold components (Fig. 2A, C-E, G)^17,18^. Side-view analysis showed that AKIP1 occupied a 272 ± 47 nm region along centrioles with an α/β-tubulin length of 468 ± 54 nm, corresponding to 57.9 ± 4.9% of the total centriole length. This signal extended from 28.0 ± 3.9% to 85.9 ± 2.7% of the centriole length when measured from the proximal end (Fig. 2B, F). This axial distribution closely matched those of POC1A, POC5, and Centrin-2/3, which covered 58.6 ± 5.3% (26.0 ± 4.1% to 84.6 ± 2.8%), 58.1 ± 3.9% (26.3 ± 3.4% to 84.4 ± 2.8%), and 57.5 ± 7.1% (25.0 ± 4.6% to 82.5 ± 5.6%; restricted to the central core-localized region) of the centriole length, respectively. POC1B showed slightly broader coverage, spanning 62.6 ± 6.9% (20.9 ± 5.3% to 83.5 ± 4.8%) of the centriole length (Fig. 2F). We then determined the radial position of AKIP1 relative to the centriolar microtubule wall by comparing the positions of maximal fluorescence intensity for AKIP1 and α/β-tubulin. AKIP1 was shifted toward the centriole lumen by 26 ± 5 nm relative to the microtubule wall (Fig. 2H). This radial position was farther from the microtubule wall than those of POC1A (18 ± 4 nm) and POC1B (11 ± 5 nm), but similar to those of POC5 (28 ± 4 nm) and Centrin-2/3 (27 ± 4.3 nm) (Fig. 2H). Together, these observations indicate that AKIP1 is a luminally positioned component of the centriole inner scaffold.

**Figure 2.**
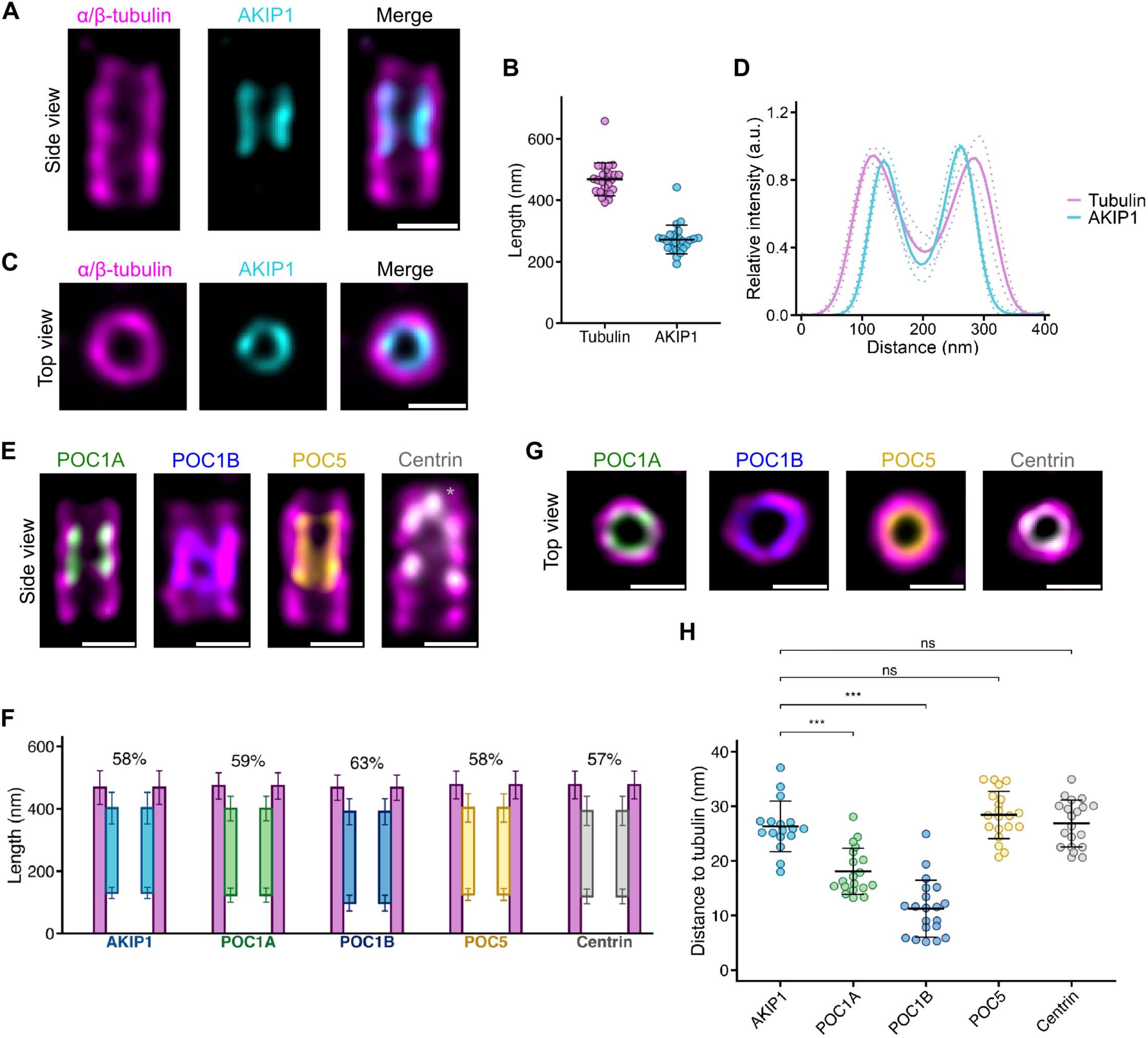
AKIP1 localizes to the inner scaffold. **(A, C)** Confocal microscopy images of RPE-1 mother centrioles expanded using U-ExM and stained for α/β-tubulin (magenta) and AKIP1 (cyan). Side view in (A) and top view in (C). Scale bar, 200 nm. **(B)** Respective lengths of α/β-tubulin and AKIP1 based on (A). Error bars, SD. n = 25 centrioles from two independent experiments. α/β-tubulin: 468 nm ± 54, AKIP1: 272 nm ± 47. **(D)** The immunofluorescence intensity profile across the two centriolar microtubules for α/β-tubulin (magenta) and AKIP1 (cyan). SDs are shown as dashed lines. **(E, G)** Confocal microscopy images of RPE-1 mother centrioles expanded using U-ExM and stained for α/β-tubulin (magenta) and POC1A (green), POC1B (blue), POC5 (yellow), or Centrin-2/3 (white). Side view in (E) and top view in (G). **(F)** Position of each stained protein along the centriole with their respective percentages of centriole coverage. Error bars, SD. Averages and SDs are as follows: AKIP1, 28.0 ± 3.9% to 85.9 ± 2.7% of centriole length, coverage 57.9 ± 4.9%; POC1A, 26.0 ± 4.1% to 84.6 ± 2.8%, coverage 58.6 ± 5.3%; POC1B, 20.9 ± 5.3% to 83.5 ± 4.8%, coverage 62.6 ± 6.9%; POC5, 26.3 ± 3.4% to 84.4 ± 2.8%, coverage 58.1 ± 3.9%; and Centrin-2/3, 25.0 ± 4.6% to 82.5 ± 5.6%, coverage 57.5 ± 7.1%, based on (B) and S1A-D, respectively. For Centrin-2/3, the tips (indicated by asterisks in (E)) were excluded, and only the region adjacent to centriolar microtubules was quantified^45^. **(H)** Distance between the maximal intensity of α/β-tubulin and the maximal intensity of AKIP1 (cyan), POC1A (green), POC1B (blue), POC5 (yellow), or Centrin-2/3 (grey) based on (E, G). Error bars, SD. AKIP1: 26.3 nm ± 4.6 (n=17), POC1A: 18.1 nm ± 4.2 (n=19), POC1B: 11.2 nm ± 5.2 (n=20), POC5: 28.4 nm ± 4.3 (n=20), Centrin-2/3: 26.9 nm ± 4.3 (n=20). A two-tailed, unpaired Student’s t test was used to obtain the P values. ***, P < 0.001; NS, P > 0.05.

### AKIP1 interacts with POC1A, POC1B, and POC5

To define the interaction network of AKIP1 within the inner scaffold, we tested its association with POC1A, POC1B, POC5, and Centrin-2 by co-immunoprecipitation. HEK293T cells expressing FLAG-tagged AKIP1 together with HA-tagged POC1A, POC1B, or Centrin-2 were subjected to immunoprecipitation with anti-FLAG antibodies, and endogenous POC5 was analyzed in the same assay. AKIP1 co-immunoprecipitated with POC1A, POC1B, and POC5, but not with Centrin-2 (Fig. 3A-D). These results suggest that AKIP1 selectively associates with POC1A, POC1B, and POC5 among the inner scaffold proteins tested.

**Figure 3.**
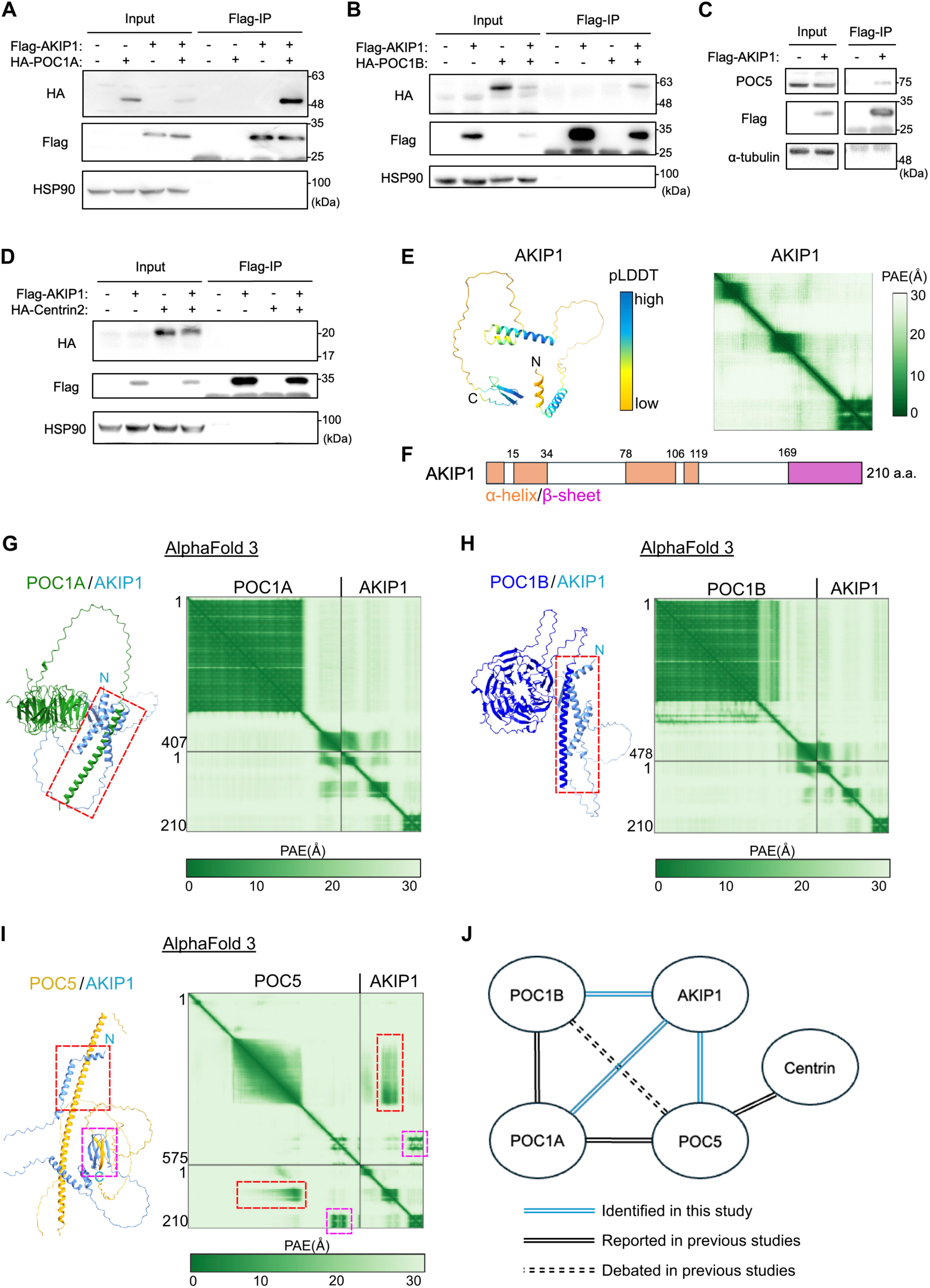
AKIP1 associates with POC1A, POC1B, and POC5. **(A-D)** Co-immunoprecipitation assay in HEK293T cells cotransfected with Flag-AKIP1 and HA-POC1A (A), Flag-AKIP1 and HA-POC1B (B), Flag-AKIP1 and HA-Centrin-2 (D), or transfected with Flag-AKIP1 alone (C). Cell lysates were immunoprecipitated with FLAG antibodies and immunoblotted with the indicated antibodies. **(E)** Predicted structure and predicted aligned error plots of AKIP1 retrieved from the AlphaFold Protein Structure Database (AF-Q9NQ31-F1), colored according to pLDDT. Blue: pLDDT > 90; cyan: 90 > pLDDT > 70; yellow: 70 > pLDDT 50; orange: pLDDT < 50. **(F)** Schematic representation of the predicted structural organization of AKIP1 based on (E). **(G-I)** AlphaFold3 predictions of complexes and predicted aligned error plots of its predictions for AKIP1 and POC1A (G), POC1B (H), or POC5 (I). Red and magenta dashed boxes indicate predicted interaction interfaces. **(J)** Schematic of the molecular interactions among known inner scaffold factors and AKIP1. Solid black lines indicate interactions established in previous studies, cyan lines represent those identified in this study, and the dashed line indicates an interaction reported previously but currently debated^17,18^.

To gain structural insight into the interactions with the inner scaffold proteins, we used AlphaFold-based protein structure predictions. First, the predicted structure of the AKIP1 monomer was retrieved from the AlphaFold Protein Structure Database^36^. This model indicated that AKIP1 contains an N-terminal α-helix, a middle α-helical region, and a C-terminal β-sheet region, corresponding to residues 1-34, 78-119, and 169-210, respectively, connected by disordered linker regions (Fig. 3E, F). We then performed complex structure predictions using AlphaFold3^37^ to model the interactions between AKIP1 and the three proteins that co-immunoprecipitated with AKIP1. AlphaFold3 predicted direct interactions between AKIP1 and each of these proteins (Fig. 3G-I). For POC1A and POC1B, their C-terminal α-helical/coiled-coil regions, corresponding to residues 357-407 and 426-478, respectively, were predicted to interact strongly with the N-terminal α-helix of AKIP1, with additional weaker contacts involving the middle α-helical region of AKIP1 (Fig. 3G, H). For POC5, an α-helical region spanning residues 155-370 was predicted to interact with the middle α-helical region of AKIP1, whereas residues 472-532 of POC5 were predicted to contact the C-terminal β-sheet region of AKIP1 (Fig. 3I). These predicted interaction modes were generally in line with additional AlphaFold-Multimer^38^ analyses and ternary-complex models of POC1A-POC5-AKIP1 (Fig. S2A-E). These results suggest that AKIP1 directly interacts with POC1A, POC1B, and POC5 through distinct structural interfaces (Fig. 3J).

### POC1A and POC5 are required for AKIP1 centriolar localization

We next examined whether the AKIP1-interacting inner scaffold components POC1A, POC1B, and POC5 are required for its centriolar localization. RNAi-mediated depletion of POC1A or POC5 in RPE-1 cells reduced the centrosomal signal intensity of the respective proteins at PCNT-positive centrosomes (Fig. S3A-D). In contrast, efficient knockdown of POC1B could not be achieved. Because POC1B has a weaker effect on the localization of other inner scaffold factors than its paralog POC1A^17^, we focused our analysis on POC1A and POC5. In POC1A- or POC5-depleted cells, the proportion of cells with reduced AKIP1 focus numbers was significantly increased compared with control cells (siControl: 0.7 ± 1.2%; siPOC1A: 81.3 ± 7.0%; siPOC5: 87.3 ± 4.2%) (Fig. 4A, B). These results indicate that the centriolar localization of AKIP1 depends on POC1A and POC5.

**Figure 4.**
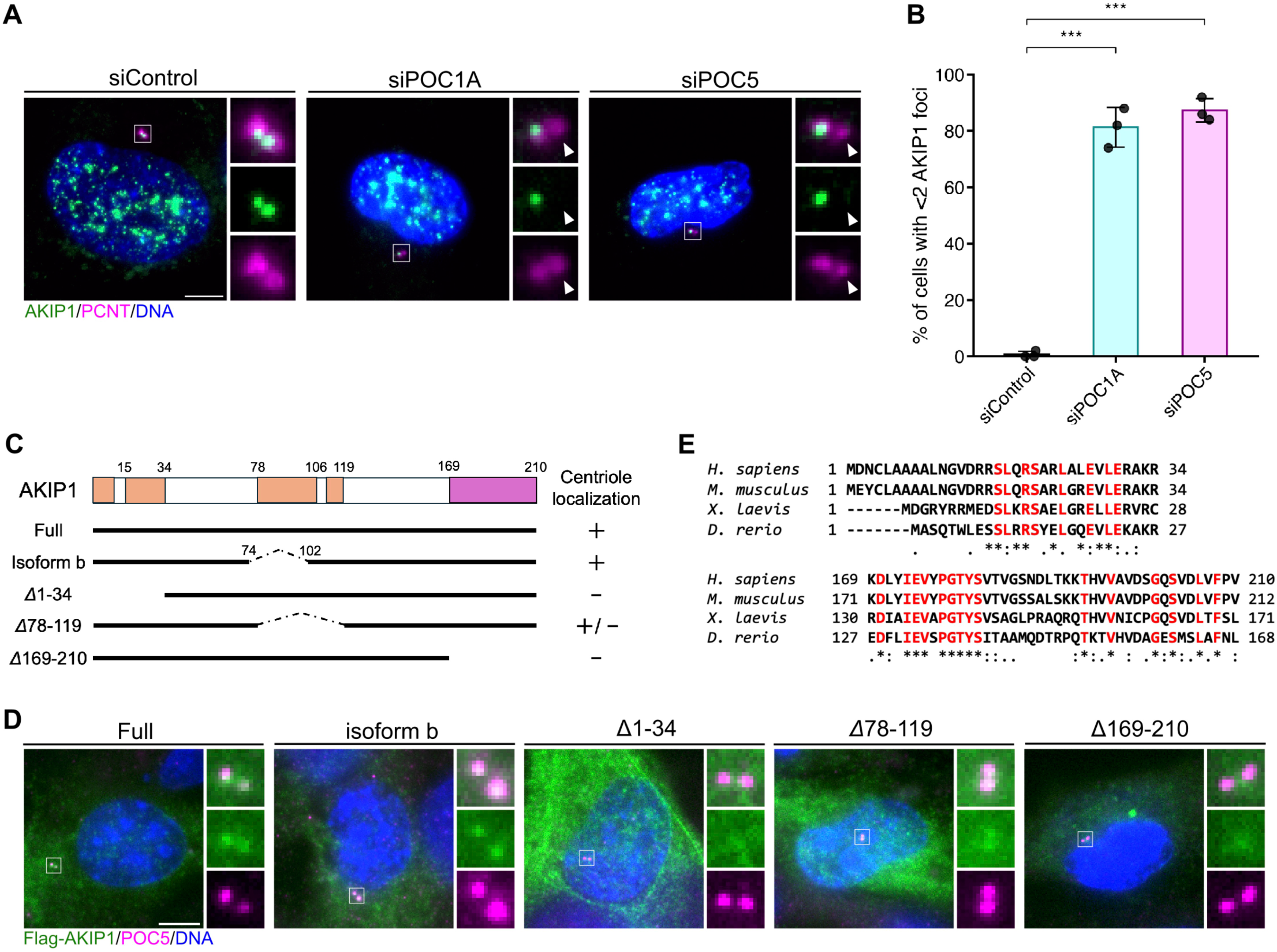
Centriolar localization of AKIP1 depends on POC1A and POC5. **(A)** Representative immunofluorescence images of control and POC1A- or POC5-depleted RPE-1 cells immunostained with antibodies against AKIP1 (green) and PCNT (magenta). DNA was visualized with DAPI. White arrows indicate no AKIP1 signal at the centrioles. Scale bar, 5 μm. **(B)** Bar graphs represent the frequency of cells with less than two AKIP1-positive centrosomes observed in (A). n = 3 independent experiments, 50 cells for each. Error bars, SD. A two-tailed, unpaired Student’s t test was used to obtain the P values. ***, P < 0.001; NS, P > 0.05. **(C)** Schematic showing specific regions of Flag-tagged AKIP1 constructs and their ability to localize to centrioles. **(D)** Representative immunofluorescence images of RPE-1 cells expressing the constructs shown in (C). RPE-1 cells were immunostained with antibodies against Flag tag (green) and POC5 (magenta). DNA was visualized with DAPI. Scale bar, 5 μm. **(E)** Amino acid sequences of the regions essential for localization identified in (C, D) in the indicated species.

We then examined which regions of AKIP1 are required for its centriolar localization by overexpressing FLAG-tagged full-length AKIP1, AKIP1 isoform b (Δ74-102), and deletion mutants in RPE-1 cells. Full-length AKIP1 and isoform b localized to POC5-marked centrioles, whereas deletion of the middle α-helical region reduced centriolar localization. In contrast, deletion of either the N-terminal α-helical region or the C-terminal β-sheet region abolished centriolar localization (Fig. 4C, D). Amino acid conservation analysis showed that these two localization-essential regions were highly conserved across species (Fig. 4E). These results indicate that the N-terminal α-helical and C-terminal β-sheet regions of AKIP1 are essential for its centriolar targeting. Together with the AlphaFold3 predictions that these regions interact with POC1A and POC5, respectively, these findings suggest that AKIP1 is likely recruited to centrioles through its conserved N-terminal α-helical and C-terminal β-sheet regions, which mediate association with POC1A and POC5, respectively.

### AKIP1 is important for maintaining centriole structure

We next investigated the role of AKIP1 in centrioles by depleting AKIP1 in cells. Because siRNA transfection did not achieve sufficient knockdown efficiency, we depleted AKIP1 by transfecting sgRNAs targeting AKIP1 into RPE-1 cells stably expressing Cas9. AKIP1-depleted cells showed a significant reduction in centrosomal AKIP1 levels compared with control cells (Fig. 5A, B). We first examined the effect of AKIP1 depletion on known inner scaffold components. Quantification of centrosomal POC1A, POC1B, POC5, and Centrin-2/3 levels revealed that AKIP1 depletion reduced POC1A levels at centrosomes, with a more modest but significant reduction in POC5 levels (Fig. 5C, D, F). In contrast, POC1B levels were not significantly altered (Fig. 5E), whereas Centrin-2/3 levels were slightly increased upon AKIP1 depletion (Fig. 5G).

**Figure 5.**
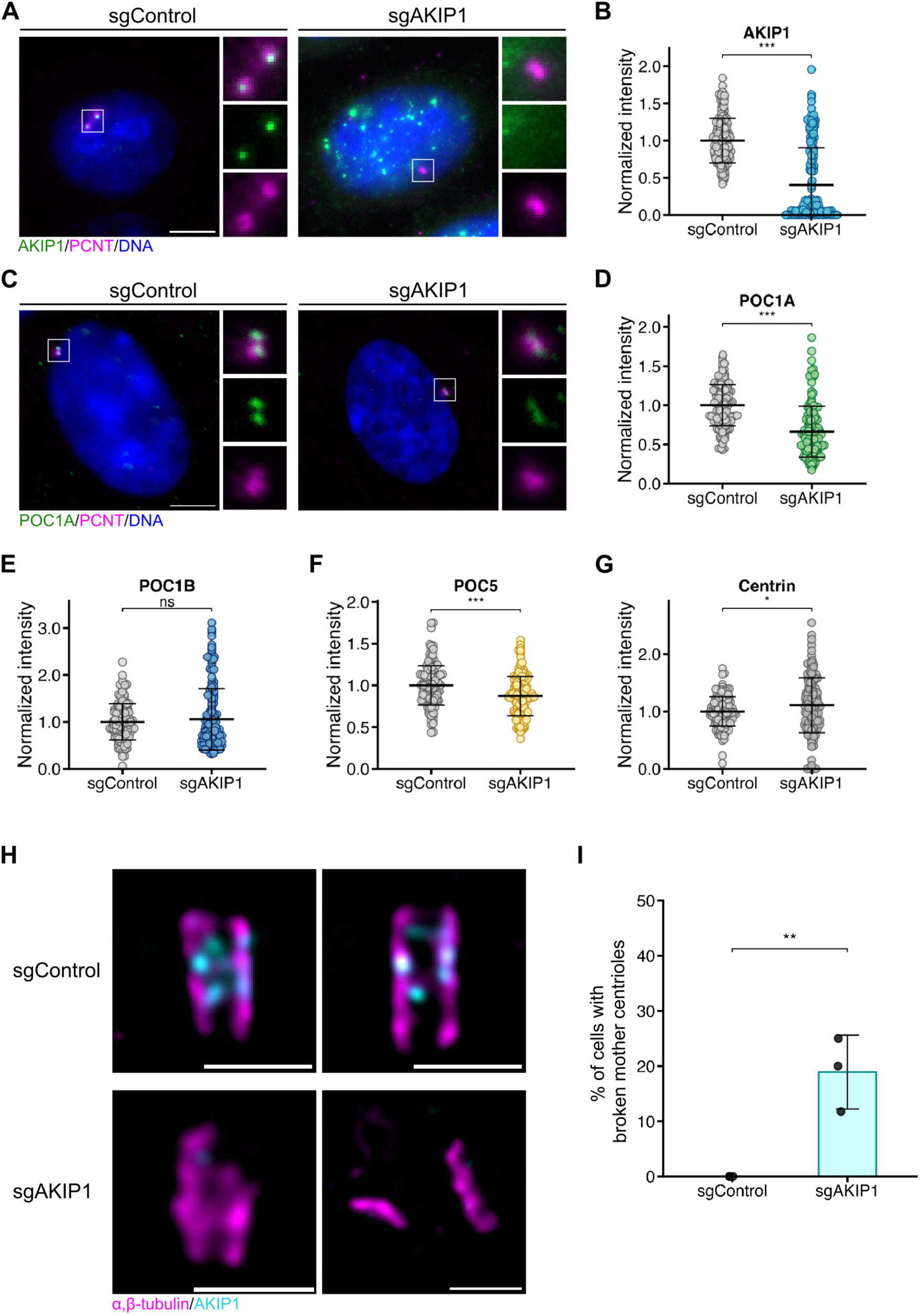
AKIP1 depletion affects the localization of other inner scaffold proteins and centriole structural integrity. **(A, C)** Representative immunofluorescence images of RPE-1 cells with CRISPR/Cas9-mediated depletion of AKIP1. RPE-1 cells stably expressing Cas9 were treated with control sgRNA (sgControl) or sgRNAs targeting AKIP1 (sgAKIP1) and immunostained with antibodies against AKIP1 (green) and PCNT (magenta) in (A), and POC1A (green) and PCNT (magenta) in (C). DNA was visualized with DAPI. Scale bar, 5 μm. **(B, D)** Quantification of the normalized signal intensity of AKIP1 in (A) and POC1A at centrosomes in (C). n = 150 cells from three independent experiments. Error bars, SD. **(E-G)** Quantification of the normalized signal intensity of POC1B (E), POC5 (F), or Centrin-2/3 (G) at centrosomes in RPE-1 cells with CRISPR/Cas9-mediated depletion of AKIP1. n = 150 cells from three independent experiments. Error bars, SD. **(H)** Representative confocal images of expanded centrioles in control and CRISPR/Cas9-mediated AKIP1-depleted RPE-1 cells stained for AKIP1 (cyan) and α/β-tubulin (magenta). Scale bar, 200 nm. **(I)** Bar graphs represent the frequency of cells with broken mother centrioles observed in (H). n = 3 independent experiments, 15-21 cells for each. Error bars, SD. A two-tailed, unpaired Student’s t test was used in B, D, E, F, G, and I to obtain P values. **, P < 0.01; *, P < 0.05; NS, P > 0.05.

Inner scaffold proteins are known to contribute to the structural stability of centrioles^14,17,21^. To determine whether AKIP1 loss affects centriole architecture, we examined centriole morphology in AKIP1-depleted cells using U-ExM. Structurally abnormal mother centrioles were observed in 19.0 ± 6.7% of AKIP1-depleted cells (Fig. 5H, I). These abnormal centrioles displayed defects such as partial loss or splitting of the microtubule wall (Fig. 5H), resembling phenotypes previously reported upon depletion of inner scaffold components^14,17,21^. These results suggest that AKIP1 contributes to the maintenance of centriole structural integrity.

Defects in some inner scaffold proteins have also been associated with impaired ciliogenesis^16,21^. We therefore examined whether AKIP1 depletion affects primary cilium formation in serum-starved RPE-1 cells. AKIP1 depletion did not significantly affect the proportion of ciliated cells compared with control cells (p = 0.588; Fig. S4A). In contrast, although the difference did not reach statistical significance, primary cilia tended to be shorter in AKIP1-depleted cells than in control cells (p = 0.0692; Fig. S4B). These results suggest that AKIP1 is largely dispensable for primary cilium formation but may have a modest effect on cilium length.

## Discussion

Although the inner scaffold supports the structural integrity and long-term stability of centrioles as a complex protein network within the centriole lumen, its full molecular composition remains incompletely defined. Here, we used large-scale gene co-dependency analysis based on DepMap genome-scale CRISPR screening data, followed by U-ExM-based validation, to identify AKIP1 as a novel component of the centriole inner scaffold. AKIP1 associates with POC1A, POC1B, and POC5, and its centriolar localization requires POC1A and POC5 as well as AKIP1 regions predicted to interact with these proteins, suggesting that AKIP1 is incorporated into the inner scaffold through these interactions. AKIP1 depletion compromises centriole structural integrity. These findings highlight gene co-dependency analysis as a useful approach for identifying components within a functional centriole network and, importantly, expand the molecular inventory of the centriole inner scaffold.

Building on these findings, several observations suggest that AKIP1 occupies a downstream position within the inner scaffold network. AKIP1 is recruited to daughter centrioles after POC5, and its centriolar localization requires POC1A and POC5, suggesting that AKIP1 is incorporated into the inner scaffold in a manner dependent on these components. The evolutionary distribution of AKIP1 may also support its downstream positioning within the inner scaffold network. Whereas the inner scaffold architecture itself is broadly conserved from *P. tetraurelia* and *C. reinhardtii* to humans, AKIP1 appears to be largely restricted to vertebrates. At the same time, co-dependency analysis suggests that AKIP1 may not be fully equivalent to the core POC1A-POC1B-POC5 module, as its correlations with these proteins were lower than those observed among the POC proteins themselves. Together, these observations suggest that AKIP1 acts as a downstream component of the inner scaffold network that contributes to centriole integrity, while also possibly retaining functional features that distinguish it from the core POC module.

The identification of AKIP1 adds a new layer to the complex protein interaction network that organizes the centriole inner scaffold. Previous work has revealed the POC1A-POC5 interaction, which is important for inner scaffold assembly and stability^17^. Our data suggest that AKIP1 can directly associate with both POC1A and POC5, and is recruited to centrioles in G2 phase, after the S-phase recruitment of POC1A^17^ and subsequent recruitment of POC5. These observations support a model in which AKIP1 is incorporated after formation of the POC1A-POC5 complex. Notably, the C-terminal region of POC5 that mediates its interaction with POC1A was also predicted to contribute to AKIP1 binding^17^, raising the possibility that AKIP1 modifies the organization of the POC1A-POC5 complex (Fig. 3I, S2A-C)^37,38^. AlphaFold3 predicted a model in which AKIP1 bridges POC1A and POC5 in place of their direct interaction, whereas AlphaFold-Multimer predicted a model in which AKIP1 binds POC5 without disrupting the POC1A-POC5 interaction(Fig. S2D, E). These models suggest two possible modes of AKIP1 incorporation: remodeling the POC1A-POC5 interaction or joining the pre-existing POC1A-POC5 complex without disrupting it. The difference between these structural models also highlights the current limitations of predicting higher-order protein complexes. Further biochemical and structural studies will be required to determine how AKIP1 is integrated into the higher-order inner scaffold network.

AKIP1 has previously been reported as a protein that binds to the catalytic subunit of protein kinase A (PKA)^35^. However, our AKIP1 co-dependency analysis did not reveal significant enrichment of PKA-related genes. This suggests that the major role of AKIP1 in cell proliferation and survival may be the maintenance of inner scaffold homeostasis. Nevertheless, because PKA is widely known to localize to centrosomes^39^, it remains possible that AKIP1 serves as one of the scaffold proteins linking PKA to centrioles. Centrosomal PKA is generally reported to localize to the PCM through AKAP9 and pericentrin^39,40^. However, some studies have shown that Centrin, an inner scaffold component, is phosphorylated by PKA, suggesting the existence of an unidentified factor that recruits PKA into the centriole lumen^41,42^. Furthermore, PKA has been reported to localize to the basal body of the primary cilium and to regulate Hedgehog signaling^43^. Future studies investigating the functional relationship between AKIP1, PKA, centrioles, and basal bodies will be important for determining whether AKIP1 provides a structural link between the centriole inner scaffold and PKA-dependent signaling pathways.

## Acknowledgments

We thank Kitagawa lab members for discussions and technical supports.

## Author contributions

Conceptualization: A.S., K.K.I., and D.K.; Methodology: A.S., K.K.I., T.C., S.H., and D.K.; Formal analysis: A.S.; Investigation: A.S.; Data curation: A.S.; Visualization: A.S.; Writing (original draft): A.S., D.K.; Writing (modification): K.K.I., T.C., S.Y., M.F. and S.H.; Supervision: D.K.; Project administration: A.S. and D.K.; Funding acquisition: S.Y., T.C., M.F., S.H. and D.K.

## Declaration of generative AI and AI-assisted technologies in the writing process

The authors used GPT-4o and Gemini 3 Pro to assist with language editing and R code refinement. Authors reviewed all suggestions provided by these tools, made necessary edits, and take full responsibility for the content of the publication.

## Material and methods

### Cell culture

RPE-1 cells were obtained from the ATCC (American Type Culture Collection), and HEK293T cells were obtained from the ECACC (European Collection of Authenticated Cell Cultures). All cells were regularly confirmed not to be contaminated with mycoplasma. The HEK293T cells were cultured in Dulbecco’s modified Eagle’s medium (DMEM; Nacalai Tesque, 08459-64) supplemented with 10% fetal bovine serum (FBS; NICHIREI, 175012), 100 U/ml penicillin, and 100 μg/ml streptomycin (Nacalai Tesque, 09367-34) at a humidified atmosphere with 5% CO₂. The RPE-1 cells were cultured in DMEM/F-12 medium (Nacalai Tesque, 11581-15) supplemented with 10% FBS, 100 U/ml penicillin, and 100 μg/ml streptomycin, at 37°C in a humidified atmosphere with 5% CO₂.

### Plasmid construction

Full-length AKIP1(CCDS7793.1) was amplified from a cDNA library of RPE-1 cells and subsequently cloned into pCMV5-FLAG. Truncated versions of AKIP1 were amplified from the full-length AKIP1 cDNA. Full-length POC1A, POC1B, and Centrin-2 cDNA was amplified from a cDNA library of RPE-1 cells. The cDNA sequences of POC1A, POC1B and Centrin-2 correspond to CCDS2846.1, CCDS31869.1, and CCDS14716.1 respectively. These constructs were cloned into pCMV5-HA. Construct amplification was carried out using KOD One PCR Master Mix (TOYOBO, KMM-201), and cloning of the constructs into vectors was performed with NEBuilder HiFi Assembly Master Mix (NEW ENGLAND BioLabs, E2621S).

### Plasmid transfection

Plasmid DNA transfection into RPE-1 cells was performed using Lipofectamine 3000 (Thermo Fisher Scientific, L30000115) according to the manufacturer’s instruction. Plasmid DNA transfection into HEK293T cells was performed using Lipofectamine 2000 (Thermo Fisher Scientific, 11668-019) according to the manufacturer’s instruction. Plasmid DNA transfection was performed 24 h before cell collection or fixation.

### RNA interference

The following siRNAs were used: Silencer Select siRNAs (Thermo Fisher Scientific) against POC1A(s226048, s24676), POC5(s43857, s43858), and Negative Control (4390843). Transfection of siRNA was performed using Lipofectamine RNAiMAX (Thermo Fisher Scientific, 13778-150). The transfected cells were analyzed 96 h after transfection with siRNA, additional siRNA was transfected 48 h after the first transfection.

### Antibodies

The following primary antibodies were used: rabbit polyclonal antibodies against AKIP1 (Atlas antibodies, HPA061391, IF 2 μg/ml, U-ExM 4μg/mL), AKIP1 (Invitrogen, PA5-106533, U-ExM 1:200), FLAG-tag (Merck, F7425, IB 1:1000), HA-tag (Abcam, ab9110, IB 1:1000), POC1A (Invitrogen, PA5-59217, IF 1:250, U-ExM 1:250), POC1B (Invitrogen, PA5-24495, IF 1:250, U-ExM 1:250), POC5 (Bethyl Laboratories, A303-341A, IF 1:1000, U-ExM 1:250, IB 1:1000), Pericentrin (Abcam, ab4448, IF 1:1000), Acetylated-Tubulin (Abcam, ab179484, IF 1:1000); mouse antibodies against FLAG-tag (Merck, F1804, IF 1:1000), HSP90 (BD Transduction Laboratories, 610418, IB:2000), Centrin (Merck, 04-1624, IF 1:1000, U-ExM 1:250), α-tubulin (Sigma-Aldrich, T5168, IB 1:1000), Pericentrin (Abcam, ab28144, IF 1:1000), CENP-F (Mitosin; BD Biosciences Pharmingen, 610768, IF 1:1000); rat antibodies against Centrin-2 (BioLegend, 698602, IF 1:500) and PCNA (Abcam, ab252848, IF 1:2000); guinea pig antibodies against α-tubulin (ABCD antibodies, AA345, U-ExM 1:1000), and β-tubulin (ABCD antibodies, AA344, U-ExM 1:1000). The following secondary antibodies were used: Alexa Fluor Plus 488 donkey anti-mouse IgG (Invitrogen, A32766, IF 1:1000), Alexa Fluor Plus 555 donkey anti-mouse IgG (Invitrogen, A32773, IF 1:1000), Alexa Fluor Plus 647 donkey anti-mouse IgG (Invitrogen, A32787, IF 1:1000, U-ExM 1:750), Alexa Fluor Plus 488 donkey anti-rabbit IgG (Invitrogen, A32790, IF 1:1000, U-ExM 1:750), Alexa Fluor Plus 555 donkey anti-rabbit IgG (Invitrogen, A32794, IF 1:1000), Alexa Fluor Plus 555 donkey anti-rat IgG (Invitrogen, A48270, IF 1:1000), Alexa Fluor Plus 647 donkey anti-rat IgG (Invitrogen, A48272, IF 1:1000), Alexa Fluor 555 goat anti-guinea pig IgG (Invitrogen, A21435, U-ExM 1:750), horseradish peroxidase-conjugated goat polyclonal antibodies against mouse IgG (Promega, W4021, IB 1:5000), and horseradish peroxidase-conjugated goat polyclonal antibodies against rabbit IgG (Promega, W4011, IB 1:5000).

### Single-guide RNA (sgRNA)

RPE-1 cells stably expressing Cas9 (RPE-1 Cas9 cells) generated in the previous study were used in this study^44^. sgRNA oligos targeting AKIP1 (#1: forward, 5′-CCGGCGTTTGTACGTTGTGT-3′; reverse, 5′-ACACAACGTACAAACGCCGG-3′, #2: forward, 5′-ACTCAGGGAGCCATGGACAA-3′; reverse, 5′-TTGTCCATGGCTCCCTGAGT-3′) were transcribed in vitro with the HiScribe T7 transcription kit (New England Biolabs, E2040L) and purified using the RNA Clean and Concentrator kit (Zymo Research, R1018). The purified sgRNAs were introduced into RPE-1 Cas9 cells using Lipofectamine RNAiMAX.

### Immunofluorescence (IF)

Cells seeded on coverslips (Matsunami, No. 1) were fixed with −20°C methanol for 7 min. After washing three times with PBS, the cells were permeabilized with PBS with 0.05% Triton X-100 (PBS-X) and subsequently blocked with 1% BSA in PBS-X for at least 10 min at room temperature. The cells were then incubated with primary antibodies overnight at 4°C and were then washed three times with PBS-X, followed by incubation with secondary antibodies for 1 h at room temperature. The cells were washed three times with PBS-X and mounted onto glass slides using ProLong Gold Antifade Mountant with DNA Stain DAPI (Invitrogen, P36935).

### Ultrastructure expansion microscopy (U-ExM)

The U-ExM protocol was performed as described previously^19^. RPE-1 cells cultured on 12 mm diameter round coverslips were incubated in 2% acrylamide + 1.4% formaldehyde diluted in PBS at 37°C for 3h or overnight. The coverslips were then placed on 35 μL of monomer solution (19% sodium acrylate, 0.1% bis-acrylamide, and 10% acrylamide) with 0.5% TEMED and 0.5% APS for 5 min at 4 °C and then for 30 min at 37 °C. The gels were then incubated in the denaturation buffer (200 mM SDS, 200 mM NaCl, 50 mM Tris pH 9) for 10 min and boiled at 95 °C for 90 min. Next, they were transferred into water at room temperature. After expansion, the gels were cut into quarters. The gels were then incubated with primary antibodies diluted in a blocking buffer (2% BSA in PBS) overnight at room temperature. After three washes with PBS, the cells were incubated with secondary antibodies in the blocking buffer for 3h at 37 °C. Expanded samples were mounted on 25mm round coverslips coated with poly-L-lysine with 10μL water and placed in an Attofluor Cell Chamber (Invitrogen, A7816) for imaging.

### Fluorescence microscopy

Immunofluorescence images of fixed cells were acquired using an Axio Imager M2 microscope (Carl Zeiss) equipped with a 63×/1.4 NA oil-immersion objective. Images were acquired as z-stacks, and maximum-intensity projections are shown. U-ExM images were acquired using a TCS SP8 microscope (Leica) equipped with a 63×/1.4 NA oil-immersion objective, and the images shown were deconvolved using Huygens Essential. U-ExM images are shown as single z-slice or z-stacks of 2,3 slices and scale bars are adjusted for the expansion factor.

### Immunoprecipitation (IP) and Immunoblotting (IB)

HEK293T cells were collected 24 h after transfection and lysed on ice in lysis buffer (50 mM Tris-HCl pH 7.5, 200 mM NaCl, 0.5% Triton X-100, 1 mM DTT, and 1:500 protease inhibitor cocktail (Nacalai Tesque, 25955-11)). The lysates were removed after centrifugation for 10 min. For IP of FLAG-tagged proteins, whole-cell lysates were incubated with M2 agarose gel conjugated with a FLAG antibody (Merck Millipore, A2220) for 2 h or overnight at 4°C. The beads were washed at least three times with lysis buffer and resuspended in SDS sample buffer (Nacalai Tesque, 09499-14) and heated for 5 min at 95 °C. Protein samples in SDS sample buffer were loaded onto 10% polyacrylamide gels, followed by transfer onto Immobilon-P membrane (Merck Millipore, IPVH85R). The membrane was blocked with 5% skimmed milk in PBS containing 0.02% Tween (PBS-T) for 30 min at room temperature. After washing three times with PBS-T, the membrane was incubated with primary antibodies in 5% BSA in PBS-T overnight at 4°C. After washing three times with PBS-T, the membrane was incubated with secondary antibodies in 5% skimmed milk in PBS-T for 1 h at room temperature. After washing three times with PBS-T, the membrane was soaked with Chemi-Lumi One L (Nacalai Tesque, 07880) before chemiluminescent detection.

### Informatics analysis

All the Chronos scores (DepMap Public 25Q3 CRISPRGeneEffect) were downloaded from the DepMap website (February 27th, 2026). The Pearson correlation coefficients between all genes were calculated. Hierarchical clustering was performed using Group Average method. A Gene Ontology (GO) enrichment analysis was performed using g.Profiler (https://biit.cs.ut.ee/gprofiler/gost) on the top 50 genes identified through co-dependency analysis (Fig. 1C).

### Protein structure prediction

The predicted monomeric structure of AKIP1 was retrieved from the AlphaFold Protein Structure Database (entry: AF-Q9NQ31-F1). Other predicted structures were obtained using both ColabFold (version 1.5.5) and the AlphaFold Server (AlphaFold 3). For ColabFold, the following settings were applied: template_mode: none, msa_mode: MMseqs2 (UniRef+Environmental), pair_mode: unpaired + paired, model_type: AlphaFold2-multimer-v2, num_recycles: 3. The AlphaFold Server was used with its default parameters. PDB files and PAE plots of the structural models ranked first among five models were selected as representatives. All representative PDB files were visualized using UCSF ChimeraX software.

### Statistics

Statistical analyses of the data were performed using R. P values were determined by the appropriate tests, described in the respective figure legends.

---

**Figure S1.**
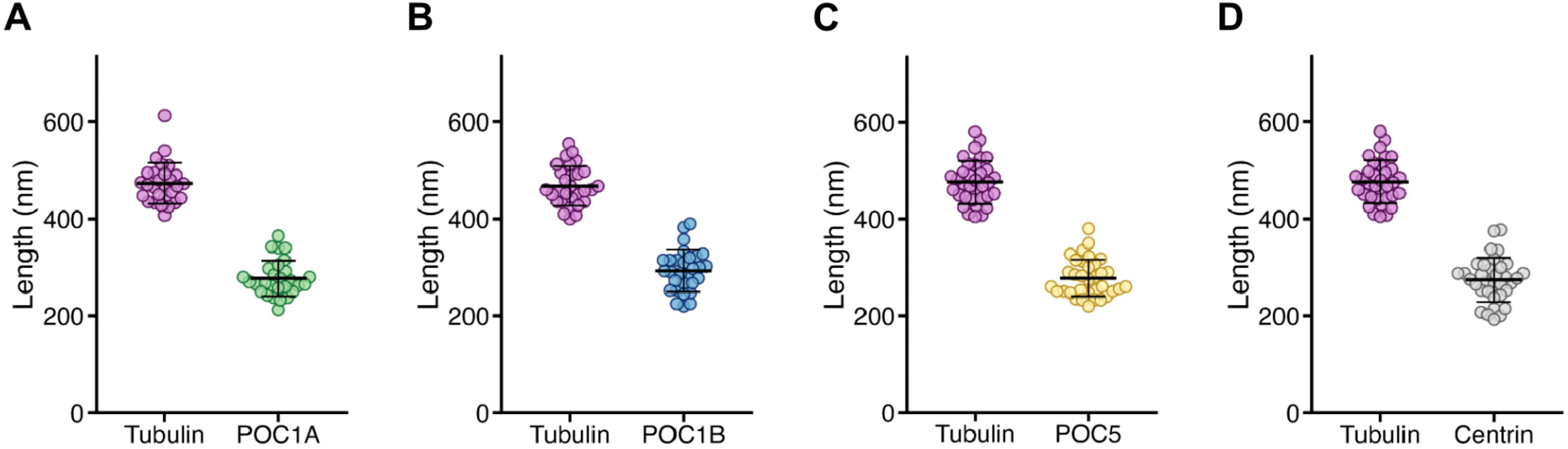
Quantification data for Figure 2F. **(A-D)** Respective lengths of α/β-tubulin and POC1A (A, n=28), POC1B (B, n=29), POC5 (C, n=34), or Centrin-2/3 (D, n=34) based on 2E. Error bars, SD. (A) α/β-tubulin: 473 nm ± 42, POC1A: 277 nm ± 37. (B) α/β-tubulin: 468 nm ± 41, POC1B: 293 nm ± 43. (C) α/β-tubulin: 476 nm ± 44, POC5: 278 nm ± 39. (D) α/β-tubulin: 476 nm ± 44, Centrin-2/3: 274 nm ± 46. For Centrin-2/3, the tips (indicated by asterisks in 2E) were excluded, and only the region adjacent to centriolar microtubules was quantified.

**Figure S2.**
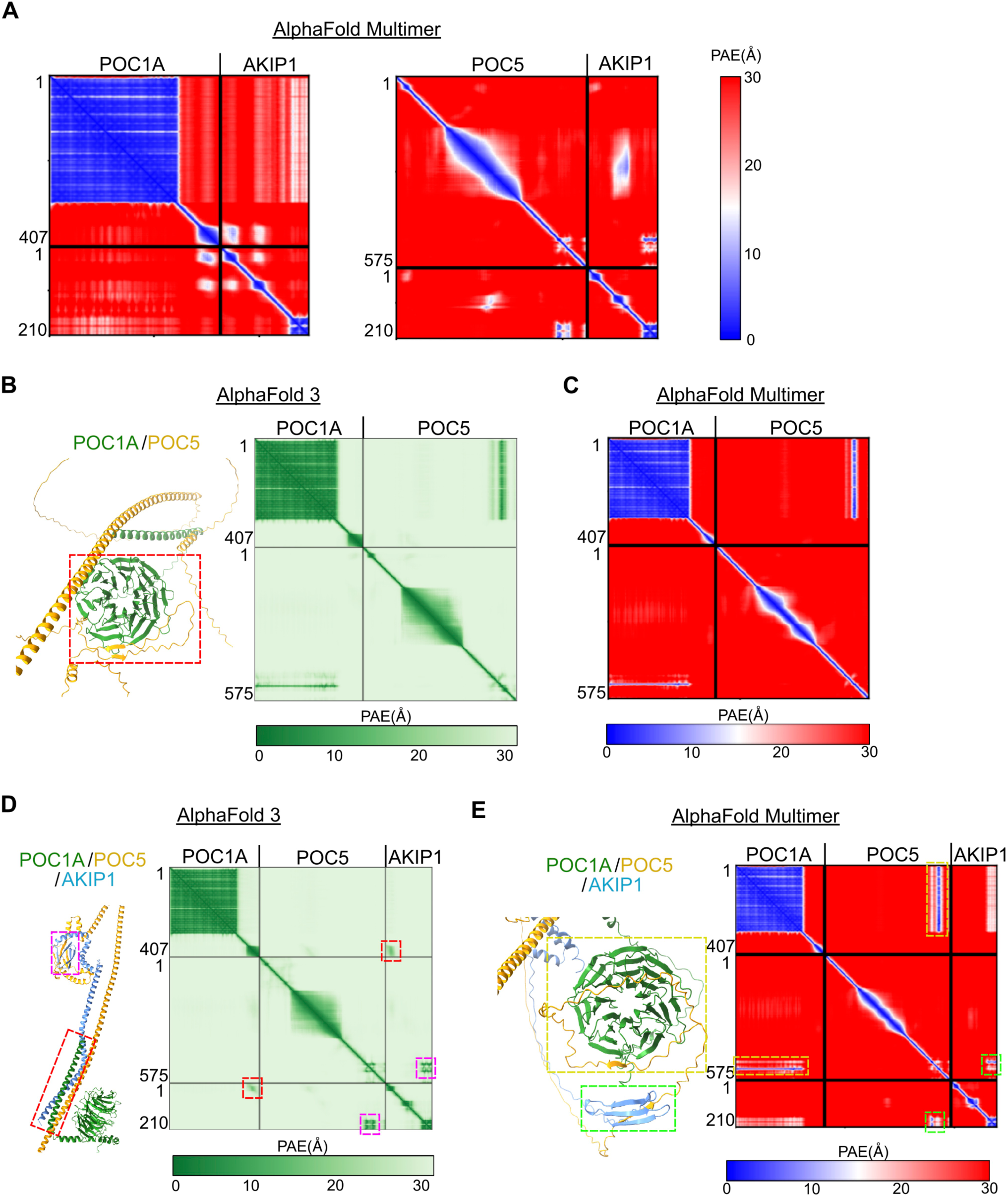
AlphaFold predictions of the ternary complex formation among AKIP1, POC1A, and POC5. **(A)** Predicted aligned error plot of POC1A-AKIP1 and POC5-AKIP1 complex structures predicted by AlphaFold-Multimer. **(B)** AlphaFold3 predictions of POC1A-POC5 complexes and predicted aligned error plots of its predictions. Red dashed boxes indicate predicted interaction interfaces. **(C)** Predicted aligned error plot of POC1A-POC5 complex structure predicted by AlphaFold-Multimer. **(D, E)** Structural predictions of POC1A-POC5-AKIP1 complexes and predicted aligned error plots of their predictions by AlphaFold3 (D) and AlphaFold-Multimer (E). Red, magenta, yellow and green dashed boxes indicate predicted interaction interfaces, respectively.

**Figure S3.**
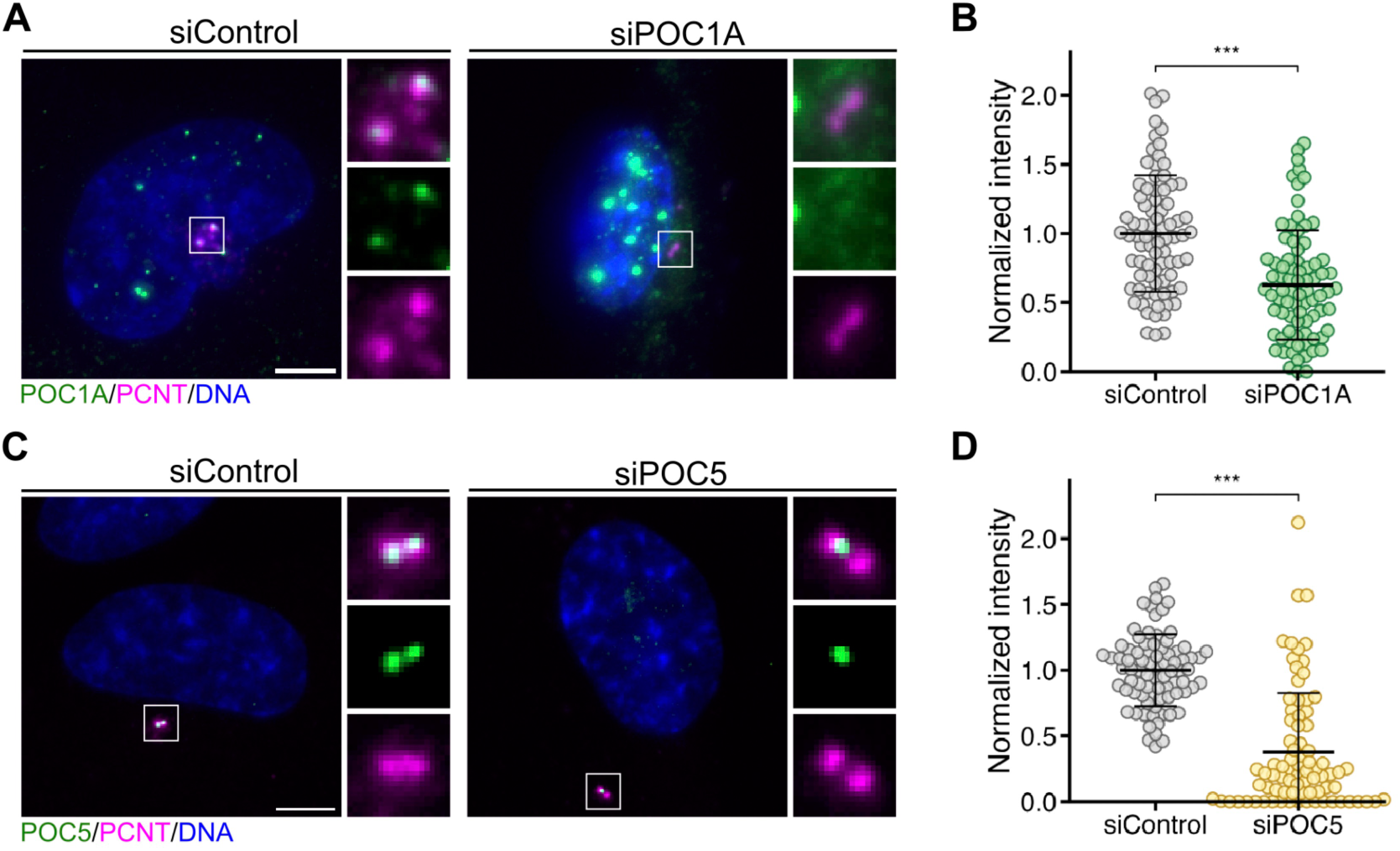
POC1A and POC5 depletion validation (for Fig. 4A, B) **(A, C)** Representative immunofluorescence images of control and POC1A- or POC5-depleted RPE-1 cells immunostained with antibodies against POC1A (green) and PCNT (magenta) in (A), and POC5 (green) and PCNT (magenta) in (C). DNA was visualized with DAPI. Scale bar, 5 μm. **(B, D)** Quantification of the normalized signal intensity of POC1A (B) or POC5 (D) at centrosomes in (A) and (C), respectively. n = 80 centrioles from two independent experiments. Error bars, SD. A two-tailed, unpaired Student’s t test was used to obtain P values. ***, P < 0.001

**Figure S4.**
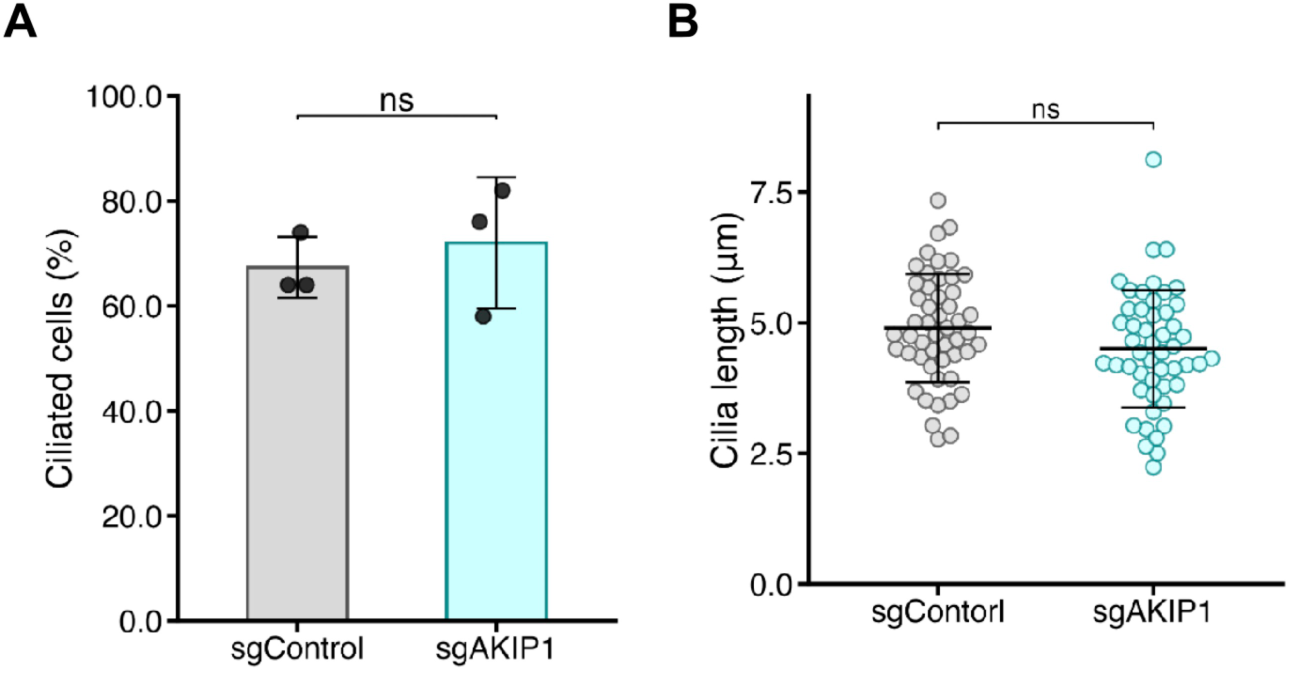
AKIP1 depletion does not affect ciliogenesis. **(A)** Bar graphs represent the frequency of cells with primary cilia. RPE-1 cells stably expressing Cas9 were treated with control sgRNA (sgControl) or sgRNAs targeting AKIP1 (sgAKIP1) and then serum starved for 48 h. n = 3 independent experiments, 50 cells each. Error bars, SD. A two-tailed, unpaired Student’s t test was used to obtain P values. NS, P > 0.05. **(B)** Quantification of length of primary cilia in the ciliated cells observed in (A). n = 50 cells from 3 independent experiments. Error bars, SD. A two-tailed, unpaired Student’s t test was used to obtain P values. NS, P > 0.05.

**Table S1.**
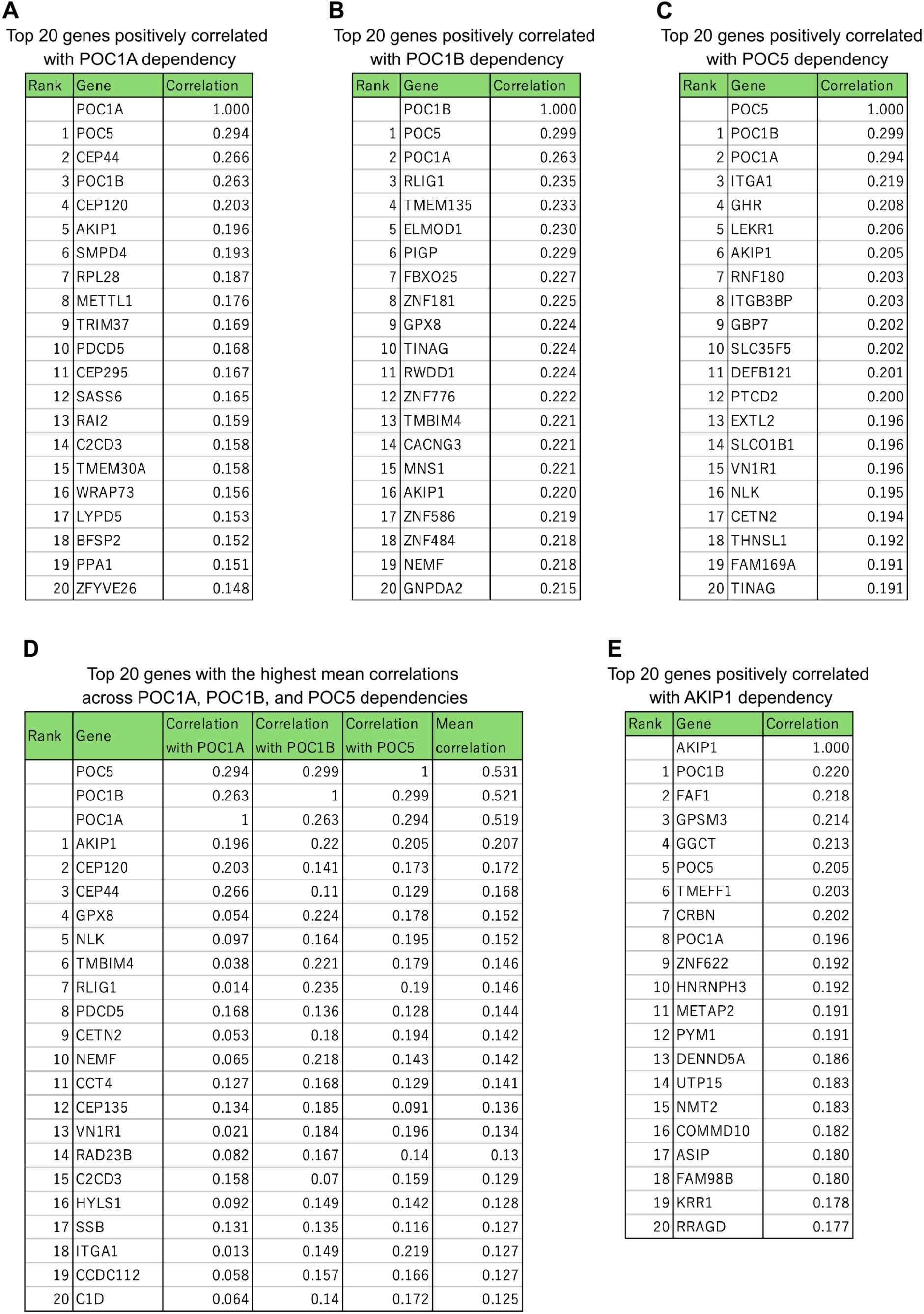
Co-dependency profiles of POC1A, POC1B, POC5, and AKIP1. **(A)** Top 20 genes positively correlated with POC1A essentiality. **(B)** Top 20 genes positively correlated with POC1B essentiality. **(C)** Top 20 genes positively correlated with POC5 essentiality. **(D)** Top 20 genes with the highest mean correlation across POC1A, POC1B, and POC5 essentialities (related to Figure 1B). **(E)** Top 20 genes positively correlated with AKIP1 essentiality (related to Figure 1D). Co-essentiality data were obtained from the DepMap Public dataset. Scores in (A)–(C) and (E) represent the Pearson correlation coefficients of CRISPR gene effect scores between each gene and the indicated target across cancer cell lines. In (D), scores represent the average of the Pearson correlation coefficients for POC1A, POC1B, and POC5.

